# Reduced regulatory complexity associated with the evolutionary transition to sociality from cockroaches to termites despite evolutionary parallels with Hymenoptera

**DOI:** 10.1101/2024.11.25.625201

**Authors:** Alun R.C. Jones, Elias Dohmen, Juliette Berger, Frédéric Legendre, Mark Harrison, Erich Bornberg-Bauer

## Abstract

The degree of phenotypic diversity within social species has previously been associated with more complex genetic regulation both in cis and trans contexts. Transcription factors (TFs) being key to genetic regulation, have been studied in the origins of eusociality in Hymenoptera (bees, ants and wasps) but less so in Blattodea (cockroaches and termites). Here we show that the social transition in Blattodea, from cockroaches to termites, displays similar patterns of regulatory change to those found in Hymenoptera. Specifically, by analysing 3 cockroach and 5 termite genomes, we find more TF families with relaxed selection compared to intensified selection and lineage-specific gene family expansions in termites, which has also been reported in Hymenoptera. We also find that genes under selection support neotenic caste determination. We find there are key differences in Blattodea TF regulation in comparison with Hymenoptera with contractions in TF gene families and no compensatory change in TF DNA binding motifs either in frequency or diversity in TF promoter regions. Furthermore, we show that an increase in social complexity leads to greater diversity in TF activating domains, one of the evolutionary and structural building blocks of TFs, meanwhile, DNA-binding domains, undergo very little change. This study highlights similarities in social transitions between Hymenoptera and Blattodea, with evidence of large changes in transcriptional regulation followed by lineage specific adaptations. Our results indicate that the transcriptional diversity linked to social complexity is not attributable to transcription factors, but is instead likely driven by an alternative mechanism..

**Significance statement:** It has been widely reported that increased phenotypic complexity, particularly arising from social transitions, is often accompanied by increased regulatory complexity followed by lineage-specific adaptations. While extensively studied in Hymenoptera (bees, ants, and wasps), it has been less explored in Blattodea (cockroaches and termites). By examining the evolution of transcription factors across Blattodea, we investigated the regulatory changes associated with this social transition. Our study reveals that increased social complexity in Blattodea is accompanied by a reduction in regulatory complexity, including decreases in transcription factor numbers and no compensatory change in TF promoter binding site diversity or regulation. This study highlights that social transitions in Blattodea and Hymenoptera share a common pattern of large regulatory changes followed by lineage-specific adaptations.

## Introduction

The evolution of eusocial organisms has remained an intriguing question in evolutionary biology as it signifies a major transition in organismal complexity [46]. The defining characteristics—cooperative brood care, reproductive division of labor, and overlapping generations—have been observed in a wide variety of taxa, including mammals, shrimp, and insects, such as bees, ants, wasps and beetles [11, 13, 33, 42, 66]. There are several other types of social organisation, such as sub sociality which involves parental care and gregarious behaviour in species which live in large groups of conspecifics [13, 30, 41]. The vast majority of eusocial research has focused on Hymenoptera [65], which harbors multiple origins of eusociality [60]. The Blattodea have a single origin of eusociality, the termites, but exhibit a full continuum of social behaviors with solitary, gregarious, sub social, primitively eusocial and complex eusocial clades allowing for comparative studies both within Blattodea and between other social taxa [7, 18, 42, 69]. Other interesting features of Blattodea, which sets them apart from Hymenoptera, are that termites are hemimetabolous in contrast to the holometabolous Hymenoptera. Termites also possess both diploid males and females within colonies including long lived reproductives of both sexes [78] with XY sex chromosomes [39] in contrast to the haplodiploid system giving rise to female workers in hymenopteran colonies. Blattodea, therefore are important to understand molecular mechanisms of social evolution as well as eusocial transitions because they enable comparisons of underlying molecular and genomic features and, consequently, to disentangle which of these features are more general if they are shared with other eusocial insects and which are more clade-specific.

Exploring the molecular details underlying the evolution of Blattodea has only become feasible recently, with a total of five termite genomes and three cockroach genomes having been published to date [7, 18, 27, 44, 48, 72, 80]. Remarkable insights from these species have identified mechanisms of viviparity (live bearing) and genomic changes related to social evolution. A hallmark of social insect molecular evolution is the regulation of chemical perception. Chemical perception is essential for many important tasks such as communication, identifying nest mates and their caste status or sensing food sources. Chemoreceptors in particular have been found to undergo drastic changes in the course of evolution and have been linked to the onset and evolution of social behaviour, among others. These changes include expansions in ionotropic receptors [27], in contrast to an expansion of odorant receptors found in Hymenoptera [86] although in Hymenoptera there is evidence to the contrary, suggesting that odorant evolution is independent of sociality and possible linked with loss of flight in females [21]. The expansion of two different major chemoreceptor families in Hymenoptera and Blattodea indicates that while functional changes, accompanied by changes in chemoreceptor families, can be conserved with major transitions the molecular mechanisms to achieve the change can vary. Another convergent pattern observed in both clades is alterations in methylation patterns which have associations with caste specific gene regulation [23, 26, 45], although this relationship is not necessarily consistent [22]. Specific regulatory pathways, such as the juvenille hormore pathway, which is known to be involved in caste determination, also convergently changed in both Blattodea and Hymenoptera [27, 67]. While specific pathways have been investigated, a more general understanding of regulatory changes related to social evolution has not been conducted in Blattodea.

To gain a better understanding of gene regulatory changes linked to sociality, the present study focuses specifically on transcription factors (TFs), whose DNA binding domain and secondary activating domains control gene expression [40]. TF influence in social evolution in Hymenoptera has revealed convergent patterns of enrichment of TF binding sites in regulatory DNA regions, accompanied by significant gains in particular gene families [34, 73]. In Hymenoptera, TF-specific gene regulatory changes have been identified across eusocial origins in bees and ants. The changes included the enrichment of specific TF binding sites in promoter regions [34, 73] in combination with significant expansions in particular gene families [12, 34, 73]. It remains unclear whether similar convergent patterns exist in termites. To this end, Harrison et al. (2018) identified expansions in zinc fingers in termites [27] but changes in regulatory regions were not studied.

This study aims to identify changes in transcription factor regulation associated with social evolution in Blattodea. Utilizing the published genomes of five termites and three cockroach species, we investigate alterations in TF-mediated regulation. Specifically, we examine changes in TF number, binding structure, binding targets, and the interplay between all of those. Our goal is to elucidate key patterns for complex social transitions in Blattodea. By comparing these findings to analogous patterns in Hymenoptera, we aim to create a more comprehensive understanding of the genetic regulatory changes associated with social evolution. This study aims to determine whether transcription factor regulatory changes associated with sociality are (A) present in non-Hymenopteran social transitions and (B) follow similar patterns to those observed in Hymenoptera.

## Results

### TF and Orthology identification

Sequences containing known *D. melanogaster* TF DNA binding domains were collected as *Drosophilla melanogaster* is the closest species with in-depth functional studies into transcription factor function and of a significantly higher quality genome than those in Blattodea,. All genomes used had high BUSCO completeness with scores above 90% except for *Blattella germanica* with a score of 84.1%. The cockroaches contained the following number of TFs; *Blattella germanica* 805, *Periplaneta americana* 1096 and *Diploptera punctata* 1274 TFs. The termites contain: *Coptotermes formosanus* 615, *Cryptotermes secundus* 911, *Macrotermes natalensis* 567, *Zootermopsis nevadensis* 681 and *Reticulitermes speratus* 854. TFs were also extracted from outgroups including *Frankliniella occidentalis* 741 and *Eriosoma lanigerum* 740. We therefore have 7,430 DNA bindingdomain containing sequences, which form a total of 12630 orthogroup sequences spread across 1620 orthogroups. There are more orthogroup sequences then DNA binding sequences as when a DNA binding domain emerges in one sequence the rest of the orthgroup do not have that DNA binding domain or they may have all had DNA binding domains but several lost over time.

### Selection

To see if TFs in Blattodea follow similar patterns of selection concerning social evolution as those found in Hymenoptera, we looked at branch specific relaxed and episodic diversifying selection as well as site-specific episodic diversifying selection in 45 single copy ortholog gene alignments. Amino acid alignments were run with GUIDANCE [71] and MAFFT [36] and trimmed below a confidence level of 0.9. Selection was determined by using a pvalue of < 0.05 for MEME and aBSREL and a false discovery rate of <0.05 for RELAX. By looking at the type of selection occurring and the specific genes, we can identify conserved patterns. RELAX tests for relaxed or intensified selection within a selection of branches. We tested this within the termites [83]. RELAX identified seven orthologs undergoing relaxed selection, all of which contain all Blattodea species, with no genes undergoing intensified selection. We also looked at the function of these genes in *Drosophila* is the most studied insect in terms of gene function. *Drosophila* is holometabolous while Blattodea are hemimetabolous, and along with evolutionary divergence, gene function in terms of TF target and biological process involved in may have changed in our study species compared to *Drosophila*.

Of the seven relaxed genes, four had variations in DNA binding domains. The zinc finger *Klumpfuss*, which is involved in neuroblast identification [35], includes a change in the C2H2-type zinc finger domain in *M. natalensis*. *Cytochrome c oxidase assembly factor 10* (*Cox10*), which is involved in heme biosynthesis and respiration [52], gains a homeodomain in *M. natalensis*, but does not have a DNA binding domain in other species. There is a gain of C2H2 zinc fingers in the vacuolar protein sorting protein 16B (*Vps16B*) in *B. germanica* protein but this is absent in other species. This gene is important for maintaining vacuole stability and structure [2]. three of these seven genes; *snoo*, *tango* and *klumpfuss*, have known neuro-developmental functions. *tango* is involved in regulating *breathless* which regulates trachea development and axonogenesis [75]. *sno* binds to a different TF which changes its affinity resulting in antagonism of the product of *dpp* and facilitation of Activin signaling [81]. *vrille* (*vri*), the bzip TF, also regulates *dpp* but as an enhancer. *dpp* is important for late stage wing development [57, 70].

aBSREL, which is an adaptive branch-site rel test, [74] that identified one gene with positive selection in two termite branches with six other genes with selection in one termite branch. All branches identified for positive selection were terminal branches. The orthogroup with selection in two tranches were in the *R. speratus* and *C. secundus* terminal branches is *Lim3*, a zinc finger and homeodomain transcription factor that regulates neuronal subtype, particularly motor neuron identity. Two genes, *Vps16* and ATPSynthase, only gained DNA binding domains in *B. germanica* and *P. americana*, so are not TFs in termites. *Macrotermes natalensis* has positive selection in *lmd* or *lame duck*, which is important for the specification of myoblasts [15]and in *abstrakt*, which is a DEAD box helicase, involved in processes such as axon asymmetric cell division [31]. *Macrotermes natalensis* also was observed with selection in the Zinc finger *CG7987* which is associated with negative regulation of transcription [79] while *C. formosanus* was found to have positive selection in Helix-loop-helix transcription factor *tango*.

MEME was used to identify sites undergoing site specific episodic diversifying selection and identified 22 sites of episodic and pervasive selection [55]. Five sites appear to undergo episodic diversifying selection. Three of these sites are in *Vps16* and one in *CEP104*, which is important for centriole elongation. Both orthgroups only gaining DNA binding domains in cockroaches. Episodic positive selection was identified in *UPf1*, which is involved in mediating nonsense mRNA degradation. Of the 11 orthogroups in which the 17 sites were identified with purifying selection, only 7 have DNA binding domains in Blattodea. One of them only gained a DNA binding domain in *M. natalensis* and another gained a DNA binding domain in *P. americana*. Orthogroups with DNA binding domains across Blattodea include: *tango*, *CG7897*, *CrebB*, *PSEA-binding protein*, *Inverted repeat protein* and *Lim3*. *PSEA-binding protein* is part of the small nuclear RNA activating complex which is involved in activating transcription of snRNA genes [29]. *CrebB* is a transcription factor important for controlling genes important for long-term memory formation [19]. *Inverted repeat protein* forms part of the Ku complex and is involved in double strand break repair and telomere maintenance [49].

### Gene family changes

In Hymenoptera, general patterns of social evolution show several lineage specific expansions, both in the TFs and in other genes [34, 73]. Consequently, we investigated gene family changes in Blattodea TFs. Using the orthofinder gene counts, we analysed gene family expansions and contractions with CAFE [47]. Orthogroups were only considered where a DNA binding domain was identified in Blattodea. At the origin of termites we found 28 gene family contractions and 6 gene family expansions (Figure 1). We then looked at significant gene family changes, which CAFE uses a likelihood ratio test and a pvalue of < 0.05 to determine. In termites at any of the branches within the clade, there are 75 significant gene family contractions compared to 63 expansions. This shows that the majority of gene family changes occur after the social origin and contractions dominate at the phylogenetic origin of eusocial evolution.

**Figure 1.**
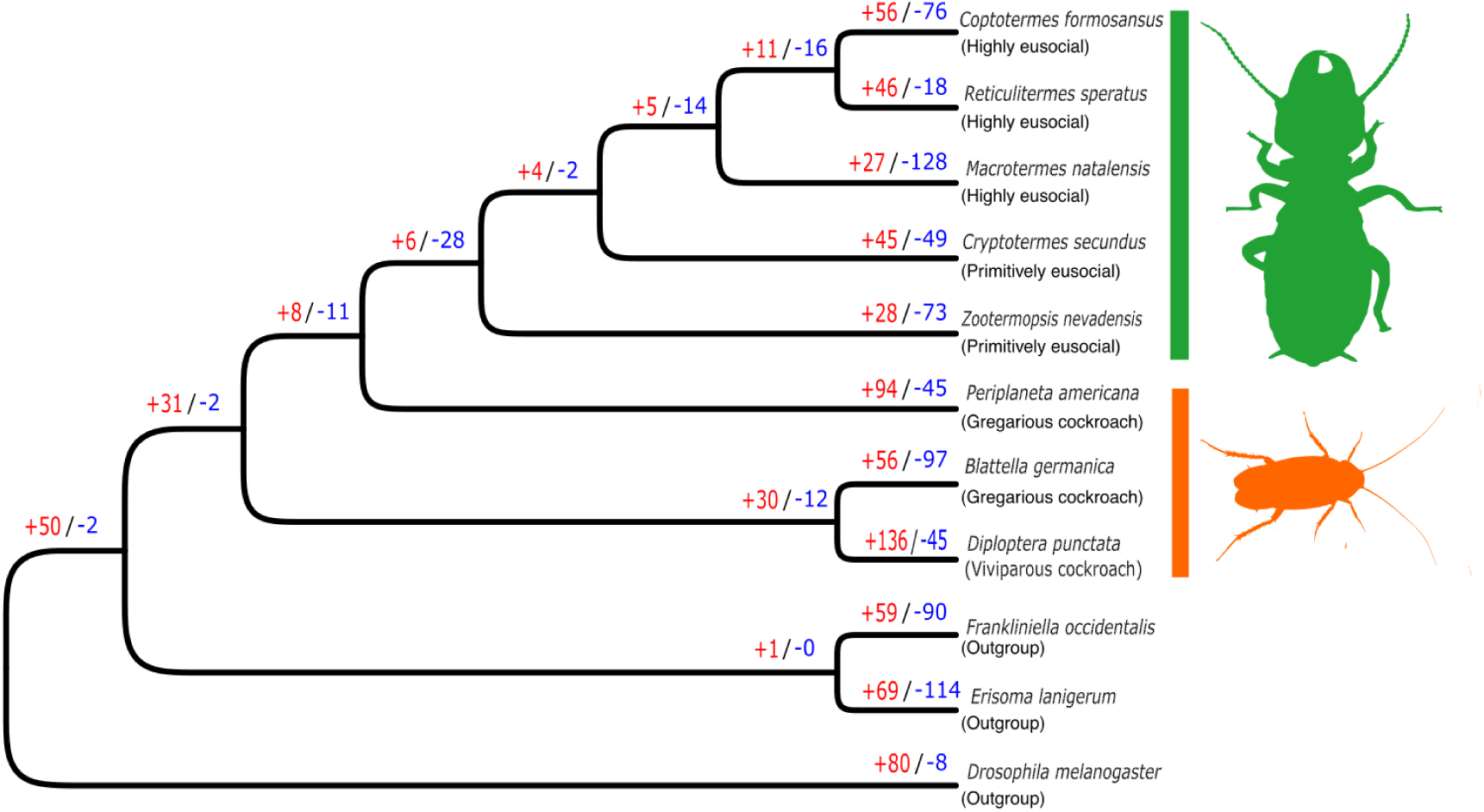
Cladogram of phylogenetic tree depicting all the species used, whether they are cockroaches or termites and their type of social organisation. Numbers in red are the number of TF containing orthogroups undergoing gene family expansions and the blue numbers are the TF containing orthogroups undergoing gene family contractions.

Given the high number of contractions, we next looked into the prevalence of contractions within the termite lineages at the terminal branches. four out of the five termite species have more contractions than expansions. Those gene families undergoing expansion are lineage-specific. There are 21 orthogroups found to only undergo contraction, while only 9 orthogroups undergo expansion. Genes families that we see undergoing expansions at the termite node include *Neuroectoderm-expressed 2*, *Rfx*, *elbow b*, *noc* and gemini (*Ubiquitin-63E*,*CG11700* but doesn’t have TF activity in fly,Ubiquitin-5E). To understand if there were similar processes affected by gene family contractions we conducted a GO term enrichment analysis. The GO term enrichment of contracted orthogroups did not show any particular function or process that was enriched.

### PFAM dynamics

To understand large scale changes in genomic regulation, it is not sufficient to just look at gene family changes but it is also necessary understand the protein function. Given that TFs are defined by DNA binding domains and activating domains, we looked at changes in the domain arrangement from our pfam scan results to understand the functional changes. Looking at the overall grouping in DNA binding domain numbers based upon the heat map there are no clear increases or decreases in DNA-binding domains among TFs that could be observed in termites (Figure 2A).

**Figure 2.**
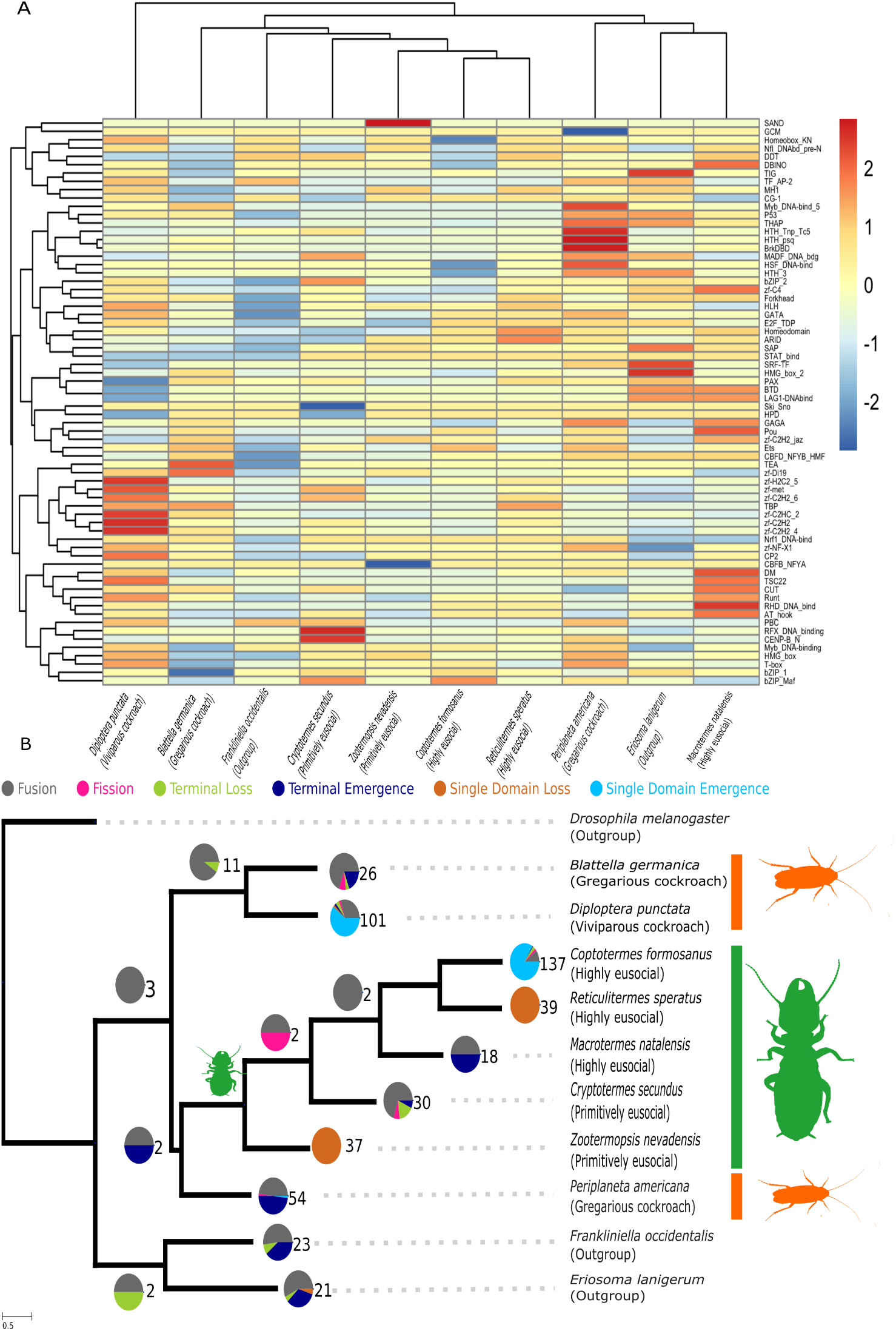
A) Heat map showing the prevalence of DNA binding domains. The number of domains has been standardised across the species representing the number of DNA binding domains within a species relative to the total mean. B) Figure depicting the output from DomRates. The numbers represent the number of domain rearrangements while the pie charts show the proportion of event types. The origin of termites is marked by the smaller green termite silhouette.

To see if there were key changes in function brought about via changes in domain modularity, we looked at protein domain rearrangements using DomRates [14]. DomRates did not show any clear patterns in TFs domain rearrangements as the type of domain rearrangement is varied in termites. Also at the emergence of the termites, there were no domain changes (Figure 2B). To understand how the protein domain and the size of the gene family relate to social complexity, we conducted a Baysian phlyogenetic generalised mixed effects model (GLMM) using the orthofinder counts and the pfam counts. We created two sets of models, one looking at the number of unique protein domains within an orthogroup and another looking at the total number of domains within an orthogroup. All model parameters converged well with Rhat values of 1 and a large Estimated Sample Size (ESS) (Sup. table 3 and 4). The full models for both sets of models had DNA binding domains and non-DNA binding domains (hereafter called activating domains) as fixed effects and a moderating effect with species social index as a factor. Orthgroup and species were included as mixed effects. Models were reduced by removing terms and all were compared using leave one out cross validation (LOO). The best-fitting model in the unique protein domains model set included a moderating effect with social index and activating domains. (Table 1). The best-fitting model for the total domain model set was the full model set (Table 2). The LOO model comparison shows that a moderating effect with social index is informative in predicting orthogroup size. We see a negative direction for the moderating affect with unique activating domains in solitary and gregarious species, while with primitively eusocial and complex eusocial species this moderating effect becomes positive (Figure 3A). A similar pattern is seen in the total domains model with the social index moderating effect that shifts the coefficients in a positive direction for the activating and DNA binding domains (Figure 3B). This indicates a small but meaningful positive relationship between the variety of domains and the total number in social species. Both sets of models show positive relationships between the number of domains and the orthogroup size meaning that several predictors have collinearity.

**Figure 3.**
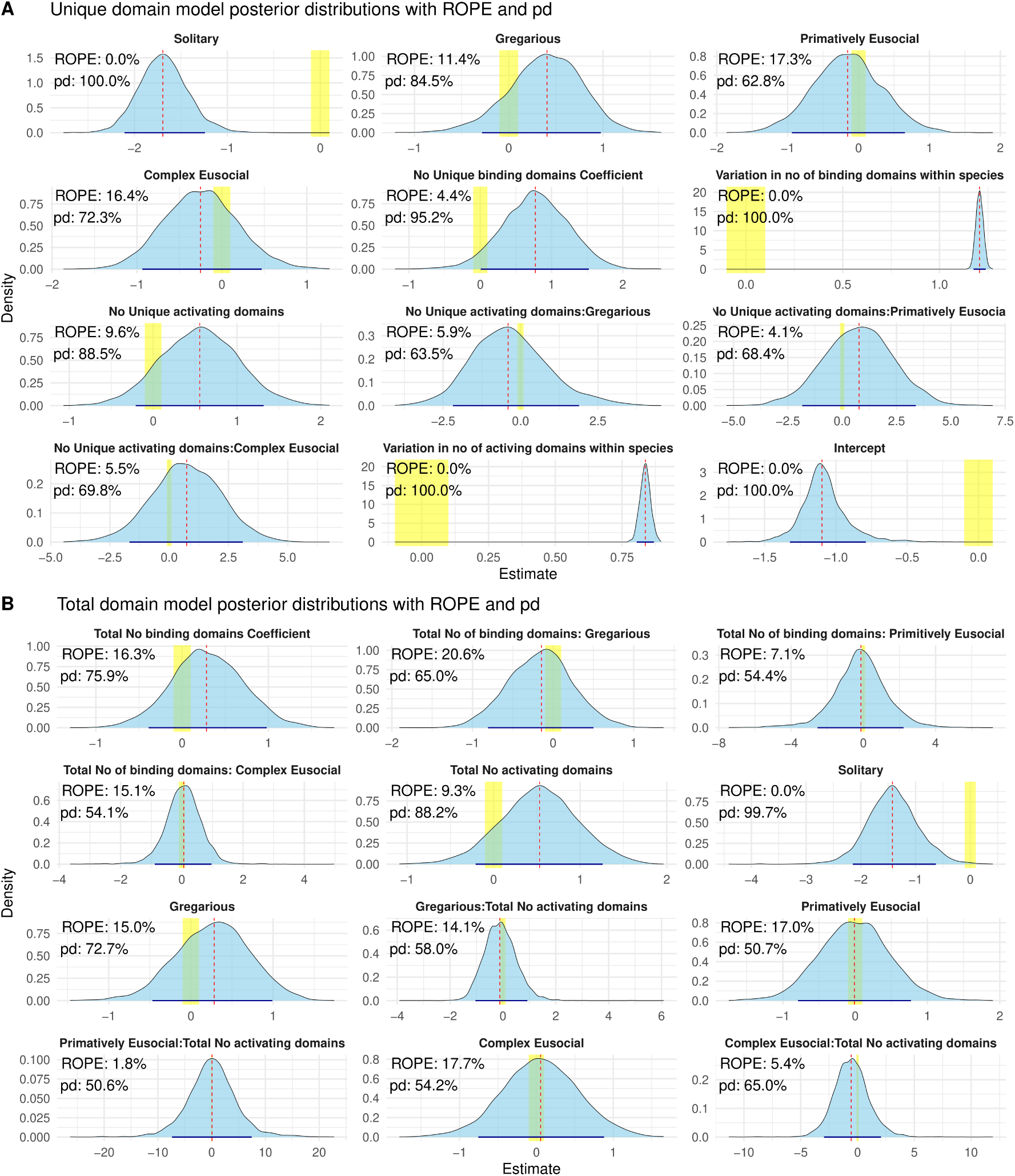
Posterior distributions of some of the coefficients of the best-fitting phylogenetic regression models, based upon LOO comparisons, predicting gene family size based upon the protein domain numbers. The yellow region is the region of posterior equivalence (ROPE) between -0.05 and 0.05 with the percentage being the percentage of the posterior distribution that lies within this region. pd percentage indicates the probability of direction (PD). The full models are in Supp table 3,4. A) Shows the best fitting model based upon the total unique number of domains within the gene family. B) Shows the best-fitting model based upon the number of unique domains within the Gene family.

### DNA binding motifs

A key finding in Hymenoptera is the enrichment in DNA binding motifs in promoter regions of social species [34, 73]. To see if the trend was present in Blattodea, we investigated DNA binding motifs in promoter regions. The promoter regions of TF genes and non-TF genes were investigated separately. DNA binding motifs were first identified using MEME and STREME [5] on the 2000bp region upstream of a start codon and only designated as a match with qvalues *<* 0.05. Motifs were checked for enrichment using SEA and deemed significant if they had a qvalue *<* 0.05 [4]. Cockroach TFs were found to be enriched with more motifs and by more diverse motifs compared to termites (Figure 4A). Given the difference in the number of TFs between the species we want to see if this is a product of differences between groups or due to the reduction in the number of TFs (Figure 1). We ran a linear model between the Shannon index and the number of promoter sequences and then looked at the difference in the residuals in a two-sided permutation test to see if there is a difference in Shannon diversity between cockroaches and termites. 10K permutations were used by randomly shuffling cockroach and termite labels and the observed difference in the Shannon diversity was -0.052 in TF promoters and 0.029 in other genes. The p-value was calculated as the proportion of permuted differences at least as extreme as the observed difference and was 0.75, indicating no significant difference between the groups in TFs and 0.69 in other genes.

**Figure 4.**
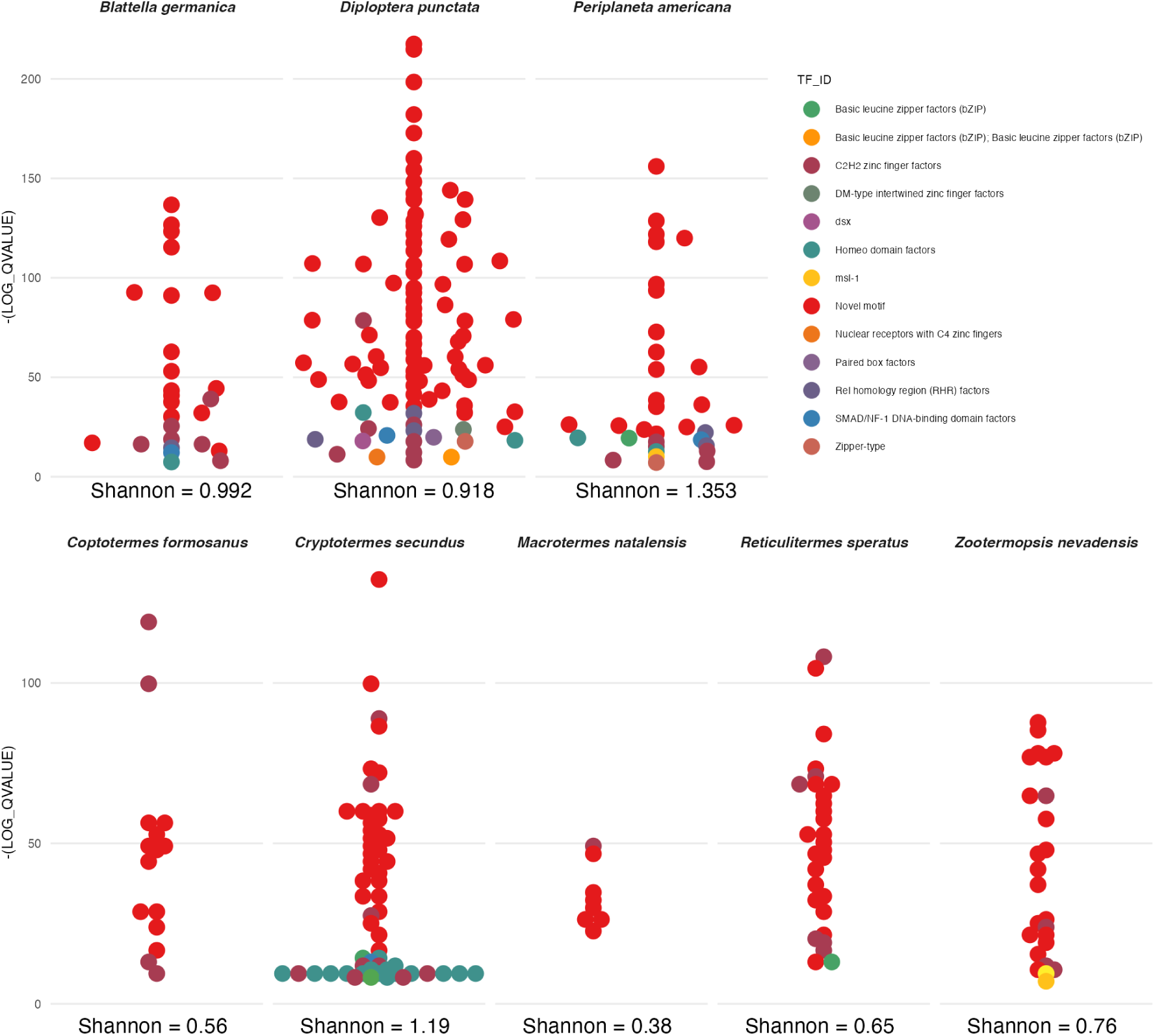
Enrichment of DNA binding domain motifs in 2kbp promoter regions of TFs in cockroaches as well as termites, identified using SEA from the XStreme package. The Shannon diversity index is displayed for each species.

## Discussion

One of the key findings of this study is that, unlike Hymenoptera, we see no real change in promoter diversity or enrichment in social termites compared to cockroaches. While Simola et al [73] and Kapheim et al [34] found that expansions in motif binding regulation are found to be correlated with higher levels of social organisation within Hymenoptera our study suggests that this is not the case for Blattodea. Increasing diversity of regulation may, therefore, not necessarily come from increasing TF binding site availability, so this would need to be considered in the context of other TF related changes, such as TF gene family changes and how the domain content within those gene families varies. Our model shows evidence that diversity is being created by changing domains, at least through the variation in activating domains (Figure 3A).

While the difference between cockroach and termite TF promoter diversity and number can be attributed to the difference in TF numbers two considerations need to be made. First, the small number of species in this study makes it difficult to rule out differences in promoters. Second, given what is known in Hymneoptera [34, 73] we predict that a reduction in TFs results in an increase in TF promoter regulation. contrastingly, we see a reduction in TFs (Figure 1) with no increase in TF regulation (Figure 4). Coupled with the overall reduction in TF families at the origin of termites (Figure 1), the results indicate increased TF regulation may not be a necessity during transitions towards eusociality. Instead, what might be more important is a general large regulatory change followed by lineage-specific changes. This trend is also observed in Hymenoptera [73], where regulation changes, such as gene family expansions, are identified alongside lineage-specific gene composition changes. Similarly, the lack of convergence in the orthogroups undergoing significant expansions in termites suggests a pattern of large changes in gene regulation followed by alterations in lineage-specific gene composition in Blattodea. At the origin of termites there are no large-scale changes in domains, but terminal branches in termites show quite a few changes in domains (Figure 2A). With the TFBS sites in the transcription factor promoter regions, while there is no significant difference between cockroaches and termites, we can see that *C. secundus* is a clear outlier compared to the other termites (Figure 4). Due to the small sample size of termite and cockroach species being studied, this relationship should be investigated in the future with a more comprehensive data set to understand if the reduction of enriched sequences is due to the loss of TFs with enriched promoters or a more general loss of binding sites. Phylogenetic models indicate weak relationships between sociality and binding domains as the social index modifying effect produces the best fitting model (Table 1,2). There is generally a positive shift in coefficients as species become more social, however despite weakly regularising priors, multi-collinearity remains a problem when interpreting these models. Larger orthogroups more protein domains than smaller ones just by having more genes in them and the number of DNA binding domains will correlate with the number of activating domains as TFs have both DNA binding domains and activating domains. The LOO model selection and ROPE results indicate that the social modifying effect is beneficial for model prediction indicating that in social species there are socially correlated changes in TF domains. This is supported by the DOMRates results (Figure 2B), as there are predominantly fusion events occurring within termites and cockroaches (Figure 2B). While major changes in domain arrangements have been found in the social transition to termites [50], this does not seem to be the case for TFs. This supports the finding in that a large change in regulation, in this case reductions rather than expansions, followed by lineage specific adaptations accompany social transitions.

When examining selection, similarities in TFs emerge between Hymenoptera and Blattodea, with relaxed selection being more prevalent than positive selection. Genes identified with both relaxed and positive selection exhibit similar functions, with many of them playing crucial roles in neuronal development and memory formation such as *tango*, *klumpfuss*, *Lim3* and *CrebB*. Notably, the only positively selected orthogroup containing a DNA binding domain in all studied Blattodea species is *Lim3*. This trend persists across multiple studies, showing consistent changes in neuronal gene expression and cell number between reproductive castes, as supported by experimental evidence [24, 62].

The global patterns found in Hymenoptera [34, 73] of relaxed selection being more prominent than intensifying selection in selection in social lineages held for Blattodea. Interestingly two genes under relaxed selection, *vri* and *sno*, are involved in regulating the *dpp* pathway which controls late stage wing development [57]. This fits in well with the neotenic workers in termites which exhibit juvenile characteristics while only alates have fully developed wings [56].

Our investigation into the regulation and composition of TFs identified that unlike expansive patterns found in Hymenoptera [73] the social transition in Blattodea is accompanied by overall reductions in TF regulation. While Hymenoptera shows an increase in transcription regulation with increased TF binding sites, Blattodea showed the opposite with fewer enriched sites, particularly in TF promoters and fewer TFs particularly losing activating domains. also identified lineage-specific changes, which is the same pattern we identify here, with social species more likely to have more unique DNA binding domains within TF containing gene families (Figure 3B). What also has to be considered, as stated earlier, is the difference in the number of species that can be investigated in Blattodea compared to Hymenoptera. It is possible that sparse sampling might show lineage-specific changes when really there are larger patterns involved. This can only be tested with more species but this study gives future studies evolutionary trends to test. This study is the first to identify a link between TF gene family diversity and sociality in termites. Future studies must look at the relationship between changes in regulatory mechanisms, instead of in isolation to understand social transitions.

## Materials and Methods

### Transcription factor identification

The defining features of TFs are the DNA binding domains and a secondary activating domain [40]. These protein domains can be identified from known modular DNA sequences allowing the identification of conserved protein functions and evolution [3, 76]. For this study transcription factors were identified using 70 Unique DNA binding PFAM domains [53] found in *D. melanogaster* transcription factors from FlyBase. We chose three cockroaches: *Blattella germanica* (84.1% BUSCO completeness), *Diploptera punctata* (97.6% BUSCO completeness) and *Periplaneta americana* (97.6% BUSCO completeness) alongside five termite species: *Zootermopsis nevadensis* (98.4% completeness), *Cryptotermes secundus* 93% BUSCO completness, *Reticulitermes speratus* (98.5% BUSCO completeness), *Coptotermes formosanus* (99.83% BUSCO completeness) and *Macrotermes natalensis* (97% BUSCO completness) additionally two outgroup species were used; *Frankliniella occidentalis* (99% BUSCO completness) and *Eriosoma lanigerum* (97% BUSCO completness) [6, 18, 27, 32, 44, 61, 68, 72, 80]. These species were chosen not only because at the time of the study they were all the genomic resources available but they also represented a large spectrum of sociality with solitary, gregarious, primitively and highly eusocial species being used. The outgroups *Eriosoma lanigerum*, the woolly apple aphid and *Frankliniella occidentalis* the western flower thrips were selected because of the quality of their genomes, their phylogenetic position to Blattodea and to keep consistency with previous studies. Pfam scans were run using the longest isoform protein sequence, and genes with any of the 70 DNA binding domains were identified as potential transcription factors. Orthogroups were identified using Orthofinder2 [16](including *D. melanogaster* 12630 TFs were identified in a total of 1621 orthogroups). Orthogroups containing genes identified as potential transcription factors were used for all analyses. Scripts for all analyses are available at https://doi.org/10.6084/m9.figshare.c.4902573.

### Multiple sequence alignment

Amino acid multiple sequence alignments of TF containing orthologs were run using GUIDANCE(2.02) [71] and MAFFT(v7.397) [36] using a global pair alignment with max iterations of 1000. Unfiltered protein alignments were transformed into nucleic acid sequences using Pal2Nal [77] and aligned but only using 200 iterations and the resulting alignments were filtered according to the amino acid alignments using a script from *Fouks et al 2021* which removed all alignments below a 90% confidence score from the final alignment [17].

### Gene family dynamics and selection

To identify TF family expansions and contractions CAFE5 [47] was used using the Orthofinder2 results. Only orthogroups where TFs had been identified in Blattodea species were used. These results were formatted using a custom bash script. A multi-lambda model was used with two lambda values estimated, one for the termite branches and one for the rest [47]. GO terms are annotated through using the PFAM2GO [54] from the sequences used in CAFE and a GO enrichment was done using TopGO with the CAFE sequences as the background universe and a significance threshold of a p-value of *<* 0.05 [1].

All 45 single-copy orthologs that were identified were used to investigate signals of selection concerning social evolution. A species tree was constructed by concatenating all single copy alignments and then running IQtree2(2.1.3) [51]. Branch-specific relaxed selection was investigated using RELAX [83], branch-specific selections using aBSREL [74] with the termites as the selected branch and sites of episodic diversifying selection using MEME [55]. Significance was determined using a pvalue of *<* 0.05 for MEME and aBSREL and a false discovery rate *<* 0.05 for RELAX. Visualisation of selection was done through Hyphy vision website.

### Protein family dynamics

PFAM domain rearrangement events were computed with DomRates [14] and visualised with the DomRates visualisation script. The total number of DNA binding domains was extracted using custom python and bash scripts and then clustered together using the R “pheatmap” package and function [37].

While genes may duplicate it does not mean that the resulting duplications have the same PFAM domain arrangements. To understand the dynamics between gene duplication and how sociality might be linked to domain changes, the relation between the number of genes in an orthogroup and the number and type of PFAM domains was modeled using a Bayesian approach. An Ornstein–Uhlenbeck (OU) covariance matrix was constructed using the species tree and the ape package vcv function [59]. Two different full multilevel negative binomial phylogenetic models were constructed using orthogroup and species as mixed effects to account for the multiple species level observations. Both full models contained fixed effects for each species’s orthogroup. One model contained the total number of DNA binding domains and the total number of activating domains while the other contained the number of unique DNA binding domains, and the number of unique activating domains. A modifier term with sociality was added for each of these fixed effects in addition to the fixed effects. Each species was given a value for the different sociality indexs: solitary, gregarious, primitively eusocial and highly eusocial. This had: *M. natalensis*, *R. speratus* and *C. formosanus* as complex eusocial and *C. secundus* and *Z. nevadensis* as primitively eusocial *B. germanica*, *P. americana* and *D. punctata* as gregarious and the outgroups as solitary. The model was run using brms(2.20.1) [9] with 4 chains on 4 cores using weakly regularising priors and a warmup of 1500 iterations and sampling of 3000. Diagnostic plots were looked at with DHarma [28]. 2 full models were reduced and compared using leave-one-out-cross-validation (LOO) [82]. Analysis was run in R(4.3.1) using the tidyverse, dplyr, ggplot and bayesplot packages [20, 64, 84, 85].

### Promoter DNA Binding

Promoter regions were extracted by taking a region of 2000bp upstream of the gene’s start codon based upon the length of known promoters [38] using GTF tools and Bedtools [8, 43, 63]. These were then split into general promoters and promoters for TFs. Using the JASPAR insect-specific TF binding motif database [10], DNA binding motifs were first identified using XSTREME which combines both MEME and STREME with default parameters [5] and only designated as a match with qvalues *<* 0.05 otherwise there were deemed ‘novel’. Motifs were checked for enrichment using SEA, which shuffles each sequence in the input set to create a control set and uses a Fisher exact test to test for enrichment against the control set. Enrichment was significant if had a qvalue of less than *<* 0.05 [4]. Motifs were compared using TOMTOM [25]. Motif comparison was done in R using JASPAR packages. The Shannon diversity index for each species was calculated using the vegan package [58]. To find if the diversity index was different between cockroaches and termites, we ran a permutation test accounting for TF number. To account for the relationship between TF number and diversity index, we ran a linear model between gene number and diversity index using Rs lm function [64]. The observed difference in residuals between cockroaches was calculated. The labels of cockroach and termite were randomly shuffled 10,000 times and a distribution of differences between group residuals was calculated. The p-value, is calculated as the proportion of permuted differences at least as extreme as the observed difference.

## Acknowledgments

ARCJ acknowledges support from DFG through grant BO 2544 / 15-1 to E.B.-B. ED was funded by the Deutsche Forschungsgemeinschaft (DFG, German Research Foundation) – 503348080. JB and FL were supported by French National Research Agency grant no ANR-19-CE02-0023 (Project SOCIOGENOMICS). Some parts of this research were conducted while visiting the Okinawa Institute of Science and Technology (OIST) through the Theoretical Sciences Visiting Program (TSVP).

## Supplementary materials

**Table 1.** Table showing the moment matched LOO comparison between phylogenetic models using the total number of unique DNA binding domains and total number of unique activating domains as predictors. Model comparison using ELPD differences, standard errors, and stacking model weights.

| Model | elpd_diff | se_diff | Stacking Weight |
| --- | --- | --- | --- |
| Unique domains model and social index modifier with activating domains | 0.0 | 0.0 | 0.683 |
| Unique full model | -6.6 | 5.5 | 0.227 |
| Unique domains model and social index modifier with DNA binding domains | -11.5 | 5.2 | 0.000 |
| Unique domains model and no modifier | -7.2 | 5.5 | 0.000 |
| Unique only DNA binding domains | -895.5 | 68.8 | 0.066 |
| Unique only activating domains | -2641.3 | 97.6 | 0.023 |

**Table 2.**
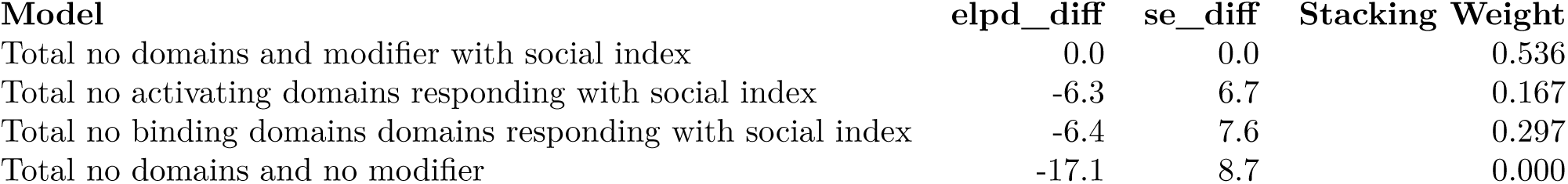
Table showing the moment matched LOO comparison between phylogenetic models using the total number of activating and DNA binding domains as a predictor. Model comparison using ELPD differences, standard errors, and stacking model weights.

**Table 3.**
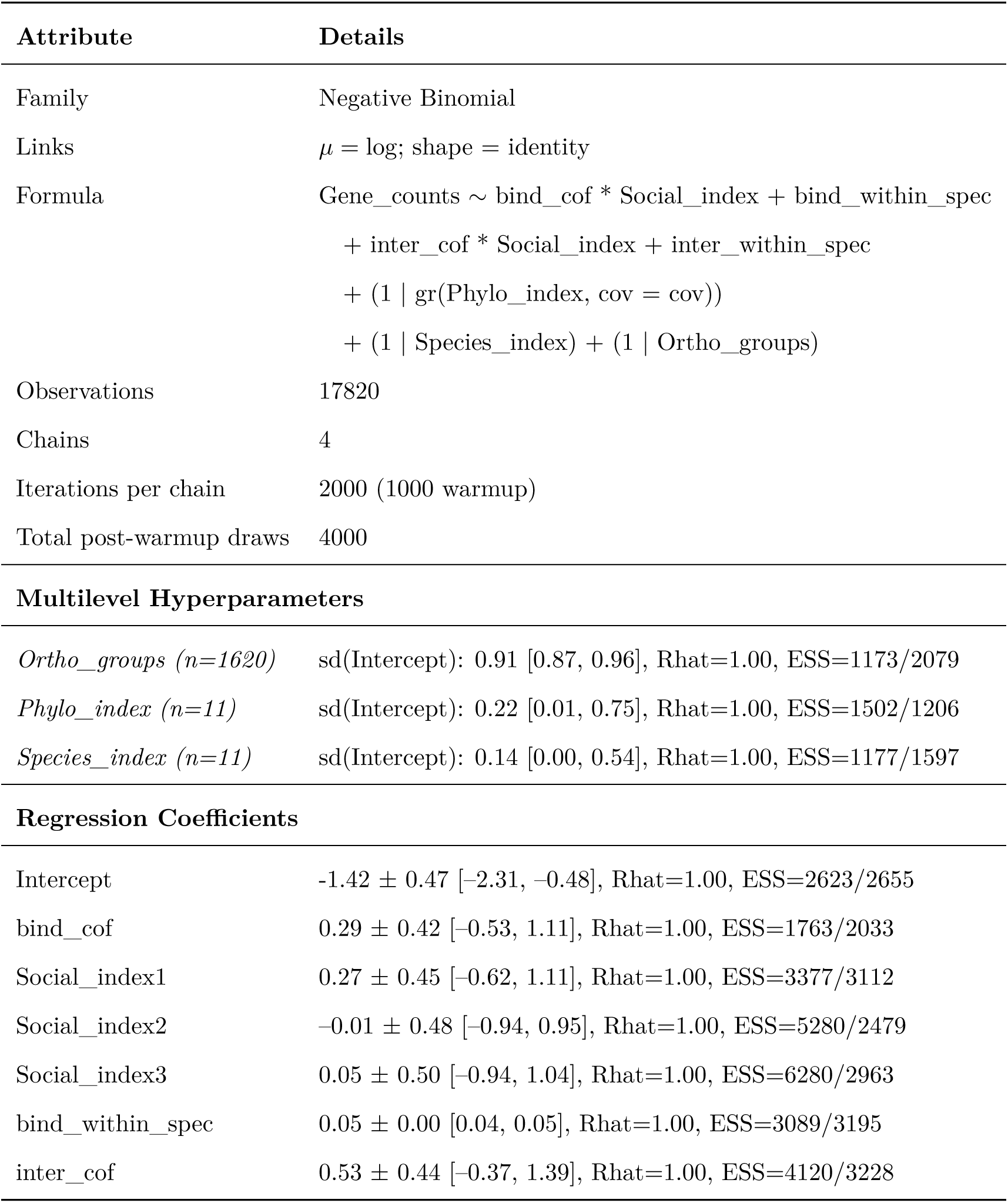
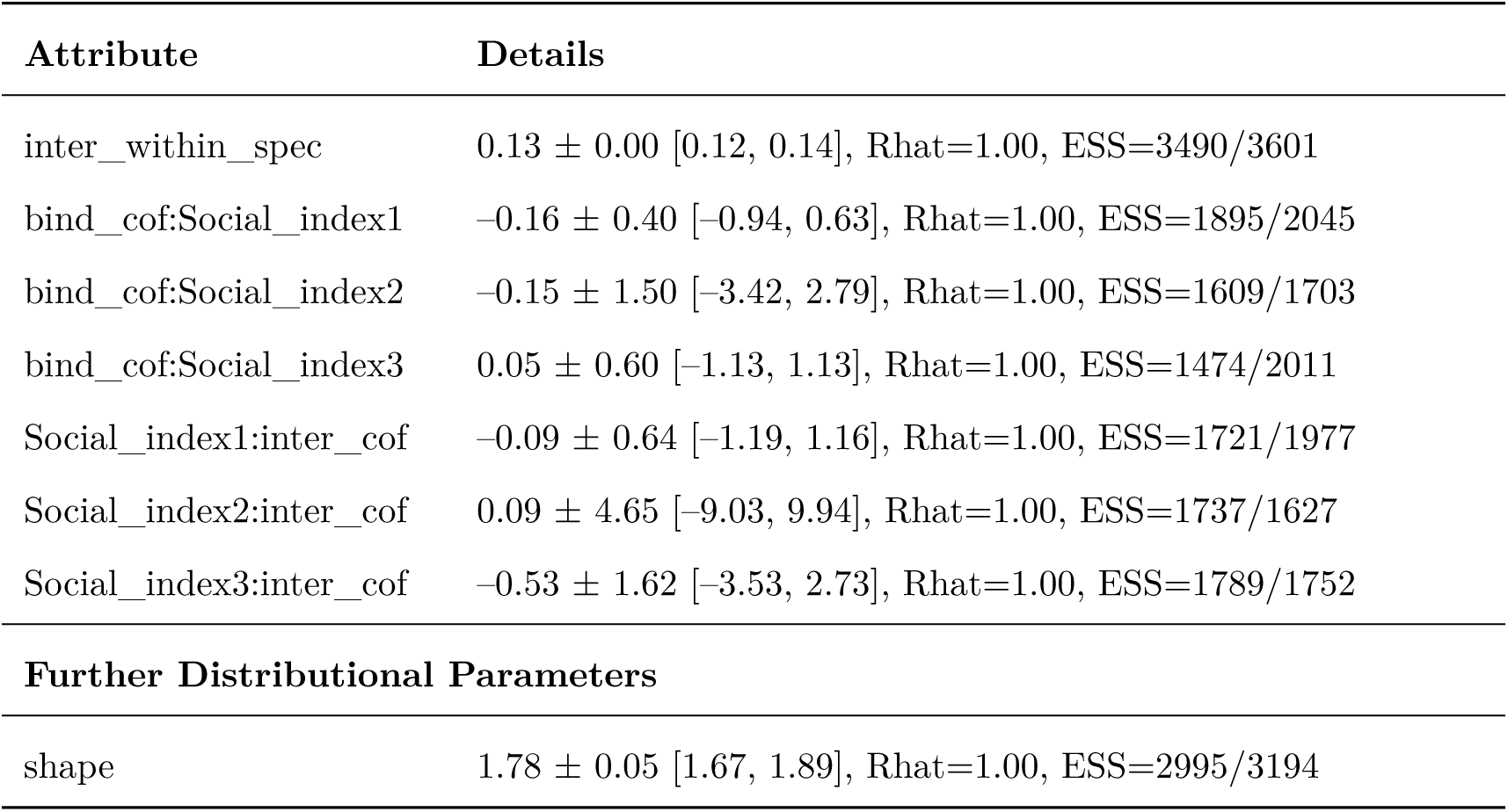
Model Summary for Total_full_model.

| Attribute | Details |
| --- | --- |
| Family | Negative Binomial |
| Links | $\mu = \text{log}$; $\text{shape} = \text{identity}$ |
| Formula | $\text{Gene\_counts} \sim \text{bind\_cof} * \text{Social\_index} + \text{bind\_within\_spec}$<br>$+ \text{inter\_cof} * \text{Social\_index} + \text{inter\_within\_spec}$<br>$+ (1 \mid \text{gr}(\text{Phylo\_index}, \text{cov} = \text{cov}))$<br>$+ (1 \mid \text{Species\_index}) + (1 \mid \text{Ortho\_groups})$ |
| Observations | 17820 |
| Chains | 4 |
| Iterations per chain | 2000 (1000 warmup) |
| Total post-warmup draws | 4000 |
| **Multilevel Hyperparameters** |  |
| *Ortho_groups* ($n=1620$) | sd(Intercept): 0.91 [0.87, 0.96], Rhat=1.00, ESS=1173/2079 |
| *Phylo_index* ($n=11$) | sd(Intercept): 0.22 [0.01, 0.75], Rhat=1.00, ESS=1502/1206 |
| *Species_index* ($n=11$) | sd(Intercept): 0.14 [0.00, 0.54], Rhat=1.00, ESS=1177/1597 |
| **Regression Coefficients** |  |
| Intercept | $-1.42 \pm 0.47$ [-2.31, -0.48], Rhat=1.00, ESS=2623/2655 |
| bind_cof | $0.29 \pm 0.42$ [-0.53, 1.11], Rhat=1.00, ESS=1763/2033 |
| Social_index1 | $0.27 \pm 0.45$ [-0.62, 1.11], Rhat=1.00, ESS=3377/3112 |
| Social_index2 | $-0.01 \pm 0.48$ [-0.94, 0.95], Rhat=1.00, ESS=5280/2479 |
| Social_index3 | $0.05 \pm 0.50$ [-0.94, 1.04], Rhat=1.00, ESS=6280/2963 |
| bind_within_spec | $0.05 \pm 0.00$ [0.04, 0.05], Rhat=1.00, ESS=3089/3195 |
| inter_cof | $0.53 \pm 0.44$ [-0.37, 1.39], Rhat=1.00, ESS=4120/3228 |
| inter_within_spec | $0.13 \pm 0.00$ [0.12, 0.14], Rhat=1.00, ESS=3490/3601 |
| bind_cof:Social_index1 | $-0.16 \pm 0.40$ [-0.94, 0.63], Rhat=1.00, ESS=1895/2045 |
| bind_cof:Social_index2 | $-0.15 \pm 1.50$ [-3.42, 2.79], Rhat=1.00, ESS=1609/1703 |
| bind_cof:Social_index3 | $0.05 \pm 0.60$ [-1.13, 1.13], Rhat=1.00, ESS=1474/2011 |
| Social_index1:inter_cof | $-0.09 \pm 0.64$ [-1.19, 1.16], Rhat=1.00, ESS=1721/1977 |
| Social_index2:inter_cof | $0.09 \pm 4.65$ [-9.03, 9.94], Rhat=1.00, ESS=1737/1627 |
| Social_index3:inter_cof | $-0.53 \pm 1.62$ [-3.53, 2.73], Rhat=1.00, ESS=1789/1752 |
| **Further Distributional Parameters** |  |
| shape | $1.78 \pm 0.05$ [1.67, 1.89], Rhat=1.00, ESS=2995/3194 |

**Table 4.**
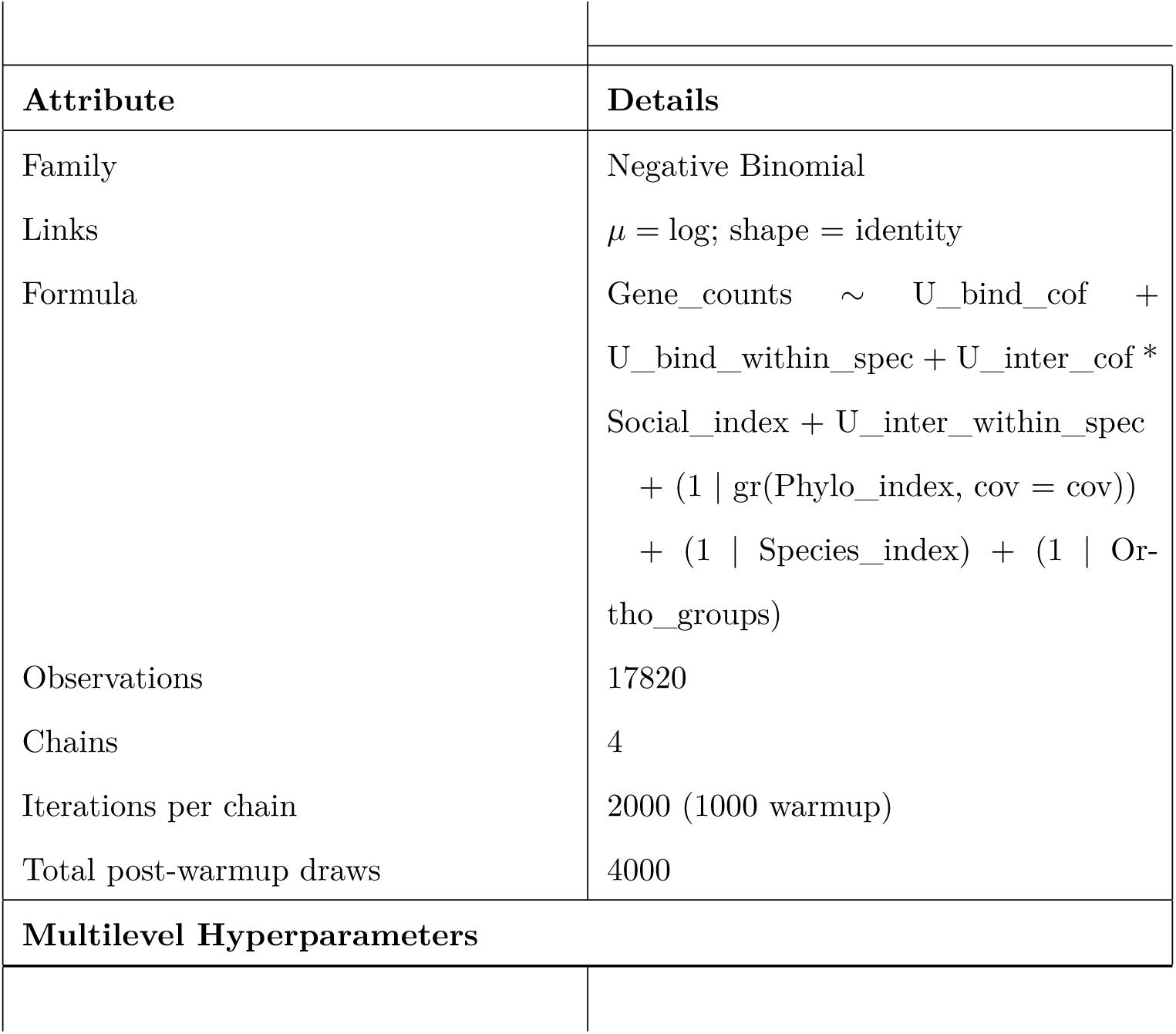
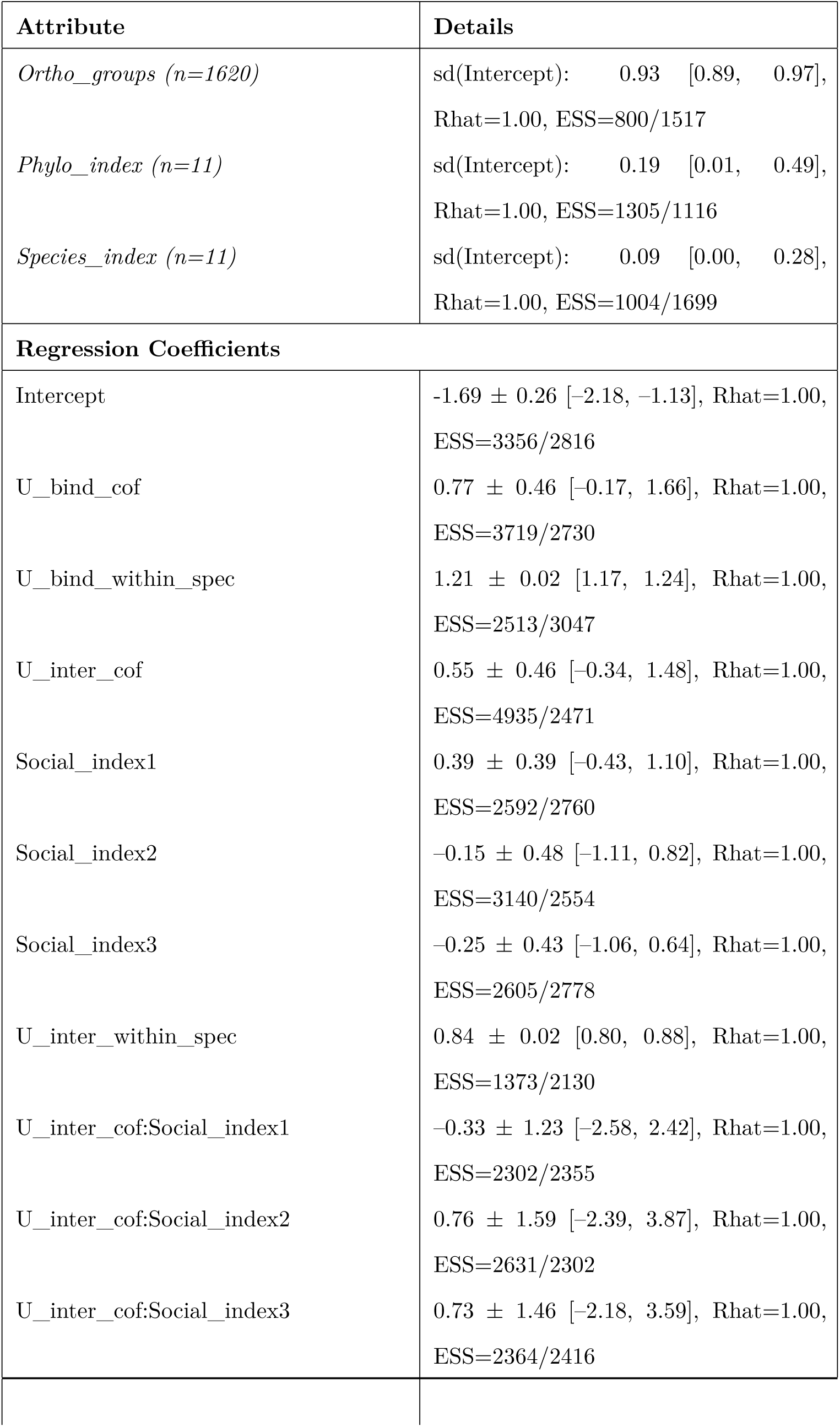
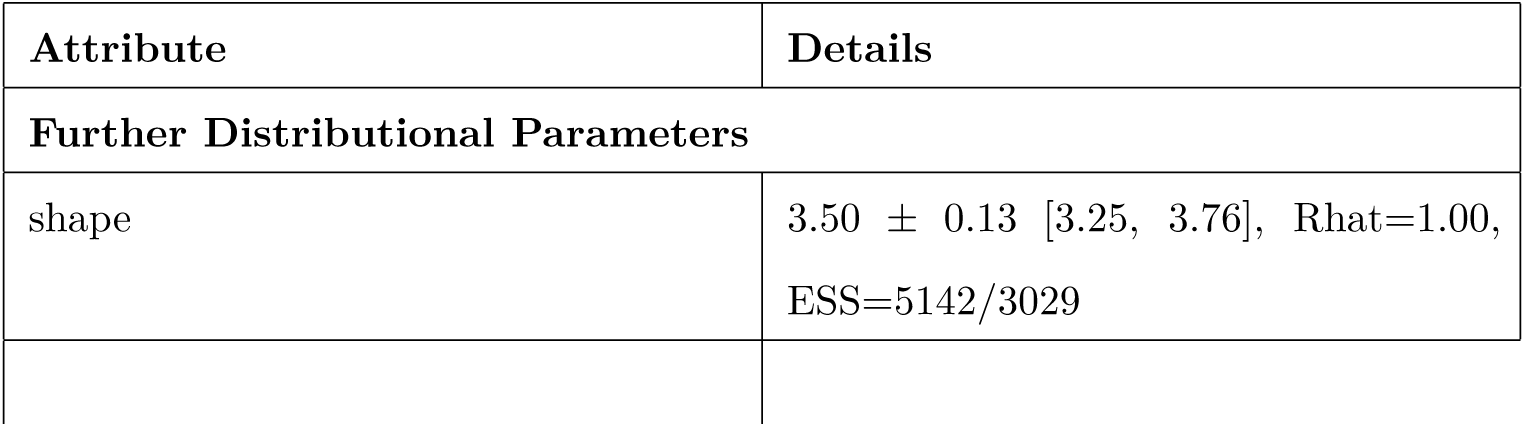
Model Summary for Unique_interacting_social_model.

